# Aberrant Wnt activation in recurrent genetically variant human pluripotent stem cells impairs cardiomyocyte differentiation and phenotype

**DOI:** 10.1101/2023.12.12.571269

**Authors:** Theodore Wing, Christopher J. Price, Dylan Stavish, Owen Laing, Jack J. Riley, Alan Lam, Steve Oh, Yaser Atlasi, Ivana Barbaric

## Abstract

Human pluripotent stem cell (hPSC)-derived cardiomyocytes have emerged as powerful tools for disease modelling and cell therapy. The production of cardiomyocytes from hPSCs typically requires expanding large numbers of hPSCs and maintaining them in culture for extended periods of time. This in turn predisposes hPSCs to the acquisition of non-random genetic changes, including recurrent gains of chromosome 1q. Here, we show that gain of chromosome 1q in hPSCs affects both the efficiency of differentiation to cardiomyocytes and phenotype of the differentiated cells. Mechanistically, we show that aberrant activation of the Wnt signalling pathway underpins the skewed differentiation of variant 1q hPSCs. Collectively, our data demonstrates that the presence of genetically variant cells in cultures is a significant concern for production of hPSC-derived cardiomyocytes for research or clinical applications. Further, our results suggest new approaches for removing genetically variant cells for future clinical applications.

## Introduction

Human pluripotent stem cells (hPSCs), which include both embryonic stem cells^1^ and induced pluripotent stem cells^2^, represent a promising source of *in vitro* derived cardiomyocytes for disease modelling and regenerative medicine^3^. Successful implementation of hPSC-derived cardiomyocytes in research and clinical practice, requires that the production of cardiomyocytes from hPSCs is reproducible and efficacious and, for cellular therapies, the resulting cells must be safe to use in humans. Concerningly, these requirements could be severely compromised by the occurrence of genetic changes in hPSCs^4^. Nevertheless, culture-acquired genetic changes are known to arise in hPSCs during expansion and prolonged maintenance of hPSCs *in vitro*^5^.

Genetic changes in hPSCs appear to be non-random with the most common changes including gains of chromosomes 1, 12, 17, 20 and X^6–8^. The non-random nature of these abnormalities is thought to result from the selective advantage they confer onto the cells under particular culture regimens^5^. Indeed, variant cells with recurrent genetic changes were previously shown to outcompete wild-type cells in culture and gain dominance through enhanced survival, decreased propensity for apoptosis, reduced cell cycle time and/or exhibiting cell competition behaviour^9–12^. Such features of variant hPSCs are reminiscent of cancer cells, raising concerns that the presence of recurrent genetic changes in hPSCs may render them unsafe to use for production of differentiated cells destined for clinical use. Another documented feature of genetically variant cells is altered differentiation propensity, which can in turn impact the efficiency and the number of produced cell types required for basic research and disease modelling. For example, previous work has shown that the amplification of chromosome 17q in hPSCs leads to altered neuronal differentiation^13^, while we recently demonstrated that the presence of isochromosome 20q impairs spontaneous differentiation of hPSCs^14^.

Although the recurrent nature of genetic aberrations in hPSCs is well established, our recent large-scale analysis of karyotyping datasets from over 23,000 hPSC cultures demonstrated that trends of karyotypic abnormalities are culture condition-dependent^15^. Specifically, we noted an increase in prevalence of chromosome 1q gains in recent years, associated with increased use of feeder-free cultures^15^. Our work^15^ and that of others^16^, showed that *MDM4* is a likely driver gene for the selective advantage of variant 1q cells, allowing variants to rapidly take over cultures through faster proliferation and reduced apoptosis. Given the prevalence of chromosome 1q variants in hPSC culture in recent years, a question arises whether the gains of chromosome 1q affect downstream application of hPSCs, such as their use for generating differentiated cells?

Here, we utilised a panel of paired lines with or without a gain of chromosome 1q to ascertain their ability to differentiate to cardiomyocytes. We show that, in comparison to wild-type cells, genetic variants with a gain of chromosome 1q exhibit impaired cardiomyocyte differentiation efficiency and altered phenotype of differentiated cells relative to WT cells. We further show that the altered differentiation of variants is mediated by aberrant activation of the Wnt signalling pathway, providing an insight into the mechanisms that are perturbed by this culture-acquired genetic aberration.

## Results

### Variants with a gain of chromosome 1q exhibit impaired differentiation to cardiomyocytes

We first set out to assess the ability of variant hPSCs with a recurrent genetic change, a gain of chromosome 1q, to differentiate into cardiomyocytes *in vitro*. To this end, we utilised three wild-type (WT) hPSC lines (H7 [WA07]^1^, H9 [WA09]^1^ and MIFF3^17^) paired with their isogenic sublines harbouring a gain of chromosome 1q (from herein termed H7 *v1q*^12^, H9 *v1q*^15^ and MIFF3 *v1q*^15^) (**Figure 1A**). For differentiation to cardiomyocytes, we used two different protocols: a commercial STEMdiff Cardiomyocyte Ventricular Differentiation kit (STEMCELL Technologies) and a microcarrier differentiation protocol adapted from previously published studies^18,19^. The commercial kit entailed differentiating cells in a monolayer by treating them with differentiation media until day 8, when the beating cell phenotype is expected to appear (**Figure 1A**). Also, following previously published microcarrier differentiation protocols, we expanded hPSCs on microcarriers for 5-7 days, prior to treating them with a Wnt pathway agonist CHIR99021 (days 0-1 of the differentiation protocol) and followed by treatment of cells with a Wnt inhibitor IWR-1 (days 3-5)^18,19^ (**Figure 1A**).

**Figure 1.**
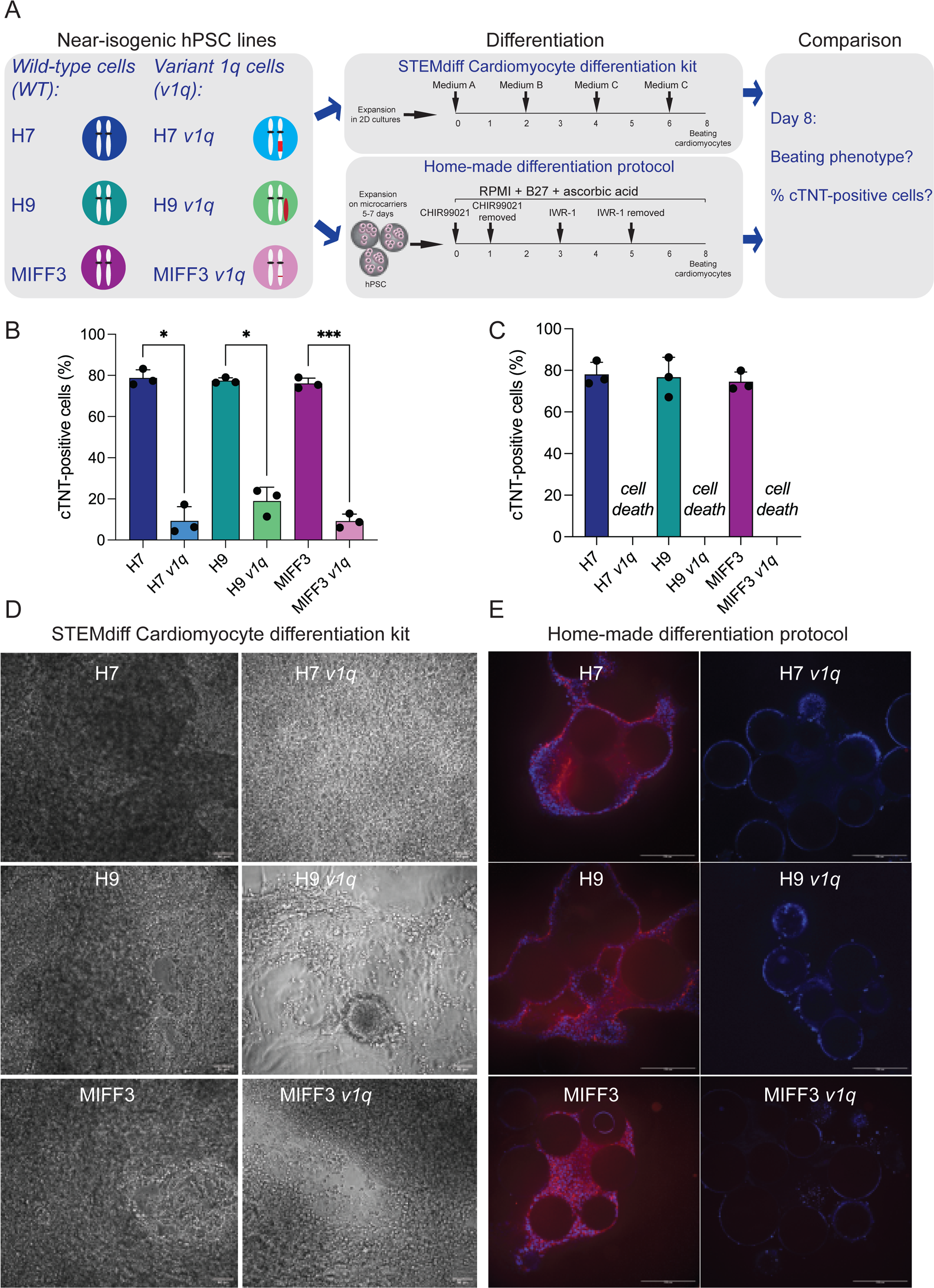
Impaired cardiomyocyte differentiation of variant hPSCs with a gain of chromosome 1q. (A) A panel of paired lines with and without a gain of chromosome 1q was investigated in two different differentiation protocols: STEMdiff Cardiomyocyte differentiation kit and a microcarrier differentiation protocol (adapted from^18,19^). At day 8, WT and *v1q* cultures were assessed for the presence of the beating phenotype and differentiation efficiency based on the percentage of cardiac troponin (cTNT) -positive cells. (B) Percentage of cTNT-positive cells for three WT lines (H7, H9 and MIFF3) and their *v1q* counterparts (H7 *v1q*, H9 *v1q* and MIFF3 *v1q*) in the STEMdiff Cardiomyocyte differentiation at day 8. (C) Percentage of cTNT-positive cells for three WT lines (H7, H9 and MIFF-3) in the microcarrier cardiomyocyte differentiation protocol at day 8. Variant cells died excessively during the differentiation protocol and could not be assessed for cTNT at day 8. (D) Representative images of WT and *v1q* cells across three different genetic backgrounds (H7, H9 and MIFF3) at day 8 of differentiation using the STEMdiff Cardiomyocyte differentiation kit. Scale bar: 50µm. (E) Representative images of WT and *v1q* cells across three different genetic backgrounds (H7, H9 and MIFF3) at day 8 of differentiation on microcarriers. Cells are stained with Phalloidin (red) and nuclei are counterstained with Hoechst 33342. Note the excessive cell death of *v1q* lines. Scale bar: 200µm. Results are the mean of three independent experiments ± SD. *p< 0.05 ; ***p< 0.001, One-way ANOVA with Šidák’s multiple comparison test. See also Figure S1A and Supplementary Videos S1-3.

As expected, at day 8 of differentiation, both protocols yielded beating cells in WT H7, H9 and MIFF3 cultures (**Supplementary Video S1, S2**). The proportion of cells positive for a cardiac marker, cardiac troponin (cTNT), was consistently above 70% for WT lines tested in each of the protocols (**Figure 1B,C)**. This was in contrast to *v1q* cells, where the differentiation efficiency was significantly reduced (**Figure 1B-E**). Neither the commercial nor the microcarrier differentiation protocol generated beating cells in variant H7*v1q* and MIFF-3*v1q* cultures at day 8 (**Figure 1D,E; Supplementary Video S3,S4**). In the H9 pair, the commercial kit generated some beating cells, but to a much lower extent than the WT ones (**Figure 1D; Supplementary Video S4)**. We observed cell death was an overt feature in *v1q* cultures in both differentiation protocols, with levels of cell death being particularly prominent in the microcarrier differentiation protocol, as less extensive *v1q* cell networks remaining attached to the microcarrier surface (**Figure 1E; Supplementary Figure 1A**). The excessive cell death of variants was surprising given the previously reported apoptotic resistance of recurrent variants under the self-renewing hPSC conditions^9–12^. Together, the differences of isogenic WT and *v1q* cells undergoing the same differentiation protocol indicated significant alterations in cell fate decisions between WT and variant hPSCs, with obvious adverse consequences for reproducibility and robustness of cardiomyocyte protocols.

### Variant 1q hPSCs are hyperresponsive to Wnt activation

To investigate the mechanisms underpinning the failure of variant cells to efficiently differentiate to cardiomyocytes, we decided to focus on utilising the microcarrier differentiation protocol as a platform for probing signalling pathways implicated in altered fate decisions in variant cells. To exclude the possibility that the increased cell death of *v1q* was due to growing the cells in a different environment (i.e. on microcarriers, rather than in conventional 2D cultures), we performed growth curve experiments of WT and *v1q* cells grown on microcarriers. We found that, much like in 2D cultures^11,12^, variant cells displayed enhanced population growth compared to WT cells during initial expansion on microcarriers (**Supplementary Figure 1B**). This suggested that the high level of variant cell death in the differentiation protocol was not due to growing the cells on microcarriers but was instead likely caused by the differentiation cues present in the medium.

The first step in our protocol for cardiomyocyte differentiation entailed activating the Wnt signalling pathway with CHIR99201. Therefore, we reasoned that the Wnt activation may be causing death and failure of variant hPSCs to differentiate. To test this hypothesis, we first treated H7 WT and *v1q* hPSCs for 18 hours in the presence or absence of 11µM CHIR99021 – the concentration for which we had already established a high differentiation efficiency in WT control lines, as shown in **Figure 1B**. As a control, we also assessed the level of apoptosis of WT and *v1q* lines grown in the E8 medium that supports the maintenance of undifferentiated hPSCs and in the RPMI medium that we used as a basal medium to induce hPSC differentiation with CHIR99021. Strikingly, while variant cells exhibited significantly lower levels of apoptosis compared to WT cells when grown in E8 or RPMI medium, they showed significantly higher levels of apoptosis upon treatment with CHIR99021 (**Figure 2A**). We noted a similar trend in H9 cells, with *v1q* sublines undergoing higher levels of apoptosis when treated with CHIR99021 compared to their WT counterparts (**Figure 2B**).

**Figure 2.**
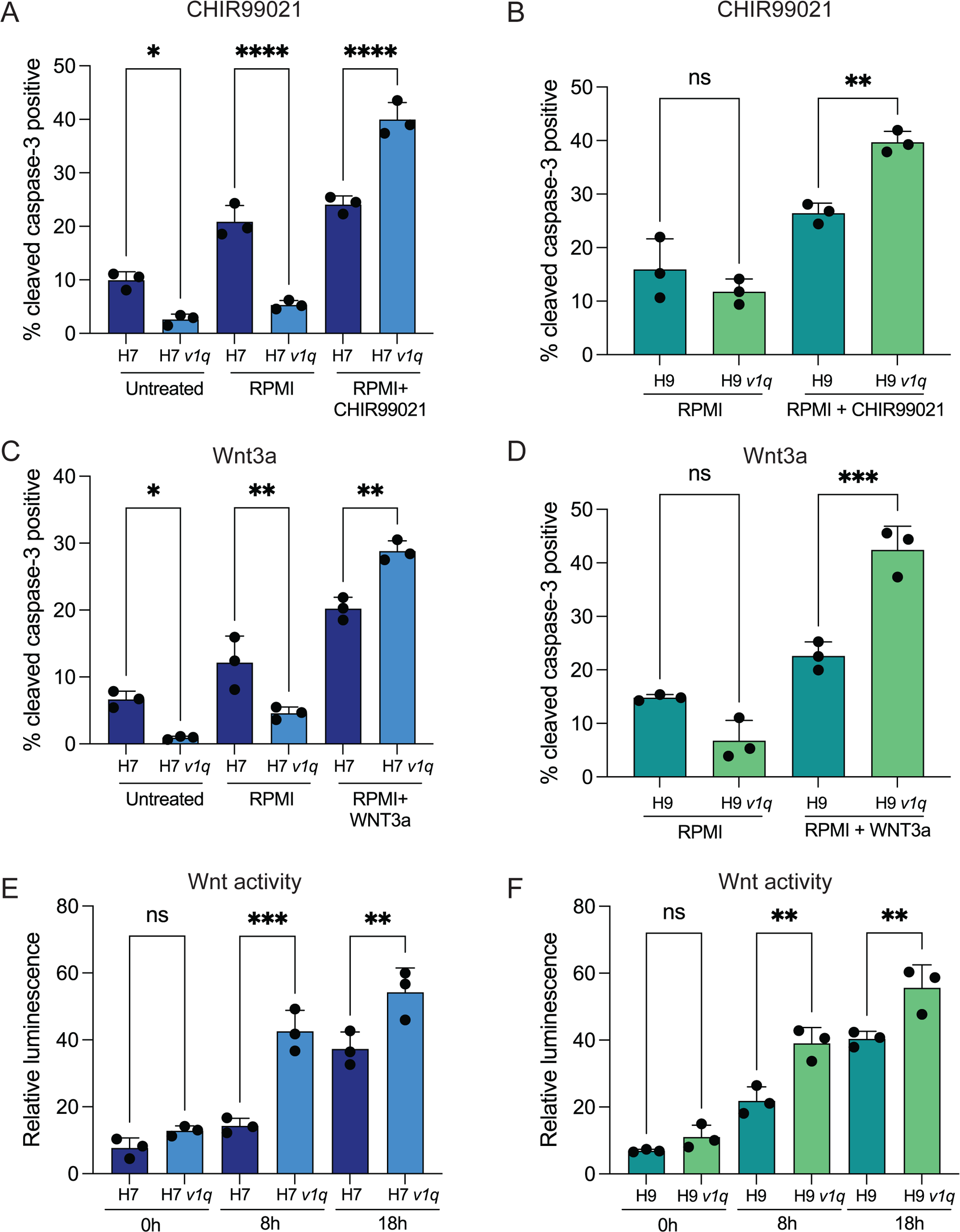
Variant 1q hPSCs undergo apoptosis upon Wnt activation. (A) Percentage of cleaved caspase-3 positive H7 and H7 *v1q* cells grown in undifferentiated state, in basal medium for differentiation (RPMI) and upon treatment with 11µM CHIR99021. (B) Percentage of caspase-3 positive cells in H9 and H9 *v1q* cells grown in basal medium for differentiation (RPMI) and upon treatment with 11µM CHIR99021. (C) Percentage of caspase-3 positive H7 and H7 *v1q* cells grown in undifferentiated state (E8), in basal medium for differentiation (RPMI) and upon treatment with Wnt3a ligand. (D) Percentage of caspase-3 positive H9 and H9 *v1q* cells grown in basal medium for differentiation (RPMI) and upon treatment with Wnt3a ligand. (E) Wnt activation measured as relative luminescence of TOPflash versus FOPflash luminescent signal in H7 and H7 *v1q* cells treated with CHIR99021 for 0h, 8h and 18h. Results are the mean of three independent experiments ± SD. *p<0.05; **p<0.01; ***p<0.001; ****p<0.0001, One-way ANOVA with Tukey’s multiple comparison test. See also Figure S1B.

Although CHIR99021 is commonly used as a Wnt agonist, this compound has multiple off-target effects^20^. Thus, to confirm that the effects observed upon the addition of CHIR99021 were specific to the stimulation of the Wnt pathway we next utilised the direct agonist of Wnt, the Wnt3a ligand, which binds to frizzled transmembrane receptors to activate downstream signalling^21^. We treated H7 WT and variant H7*v1q* cells with 200ng/ml Wnt3a ligand for the same 18 hours window and then assessed the proportion of cells expressing the cleaved caspase-3 marker of apoptosis. In a similar manner to CHIR99021 treatment, the treatment with Wnt3a caused a sharp increase in the percentage of cleaved caspase-3 positive apoptotic cells in 1q variant hPSCs compared to the WT control (**Figure 2C**). We confirmed that this result was reproducible in an additional hPSC line, H9 (**Figure 2D**). Overall, this data is consistent with the notion that Wnt activation in *v1q* hPSCs triggers predominantly apoptosis rather than differentiation of variant cells.

We next reasoned that two alternate possibilities could explain our results. One explanation is that Wnt signalling is unperturbed in variant cells, but the same level of activation leads to a different cell fate. Alternatively, the Wnt signalling may be aberrantly activated in variants, thus causing differences in cell fate between WT and *v1q* cells. To test these possibilities, we directly measured Wnt activity in WT and *v1q* cells in undifferentiated state and at different time points post-CHIR99201 treatment using the TOP-Flash Wnt reporter assay^22^. We noted no significant differences in basal Wnt signalling in WT and *v1q* cells (H7 pair) grown under conditions that support undifferentiated stem cell phenotype (0hr) (**Figure 2E**). However, an 8-hour treatment with 11µM CHIR99201 resulted in a significantly higher Wnt activation in the *v1q* subline compared to WT cells, which remained significantly elevated at the 18h timepoint (**Figure 2E**). We also noted a similar trend of Wnt activation upon CHIR99201 treatment in H9 WT and *v1q* pair lines (**Figure 2F**). Based on these experiments, we concluded that WT and *v1q* cells display significant differences in their capacity for Wnt activation, with variants responding more rapidly and achieving a higher level of Wnt activation compared to WT hPSCs.

### Impaired differentiation of *v1q* cells is caused by hyperactivation of Wnt

While the microcarrier differentiation protocol yielded no apparently beating cardiomyocytes from *v1q* cells in any of the three lines tested, the fact that we saw some cTNT-positive cells when using the STEMCELL kit (**Figure 1B,D**), as well as some beating cells in H9 *v1q* cultures suggested to us that *v1q* cells are not inherently refractory to differentiation. Based on our results that *v1q* cells are hyperresponsive to Wnt activation, we postulated that overactivation of Wnt is preventing *v1q* cells from differentiating. Hence, we hypothesized that reducing the levels of Wnt agonism in *v1q* cells would reduce the amount of cell death and permit differentiation to cardiomyocytes.

In line with our hypothesis, treatment of variant cells with lower doses of CHIR99201 (4µM) for 18h induced similar level of apoptosis in *v1q* cultures compared with their WT counterparts (**Figure 3A-C**). Notably, lowering the concentration of CHIR99201 was also effective in overcoming the differentiation block in *v1q* sublines, as all the *v1q* sublines produced beating cardiomyocytes after 8 days of the differentiation protocol (**Figure 3D**; **Supplementary Video S5).** Based on the proportion of cTNT positive cells, the efficiency of differentiation of variant cells treated with lower concentrations of CHIR99201 was more similar to those of WT cells in the original protocol (**Figure 3E**).

**Figure 3.**
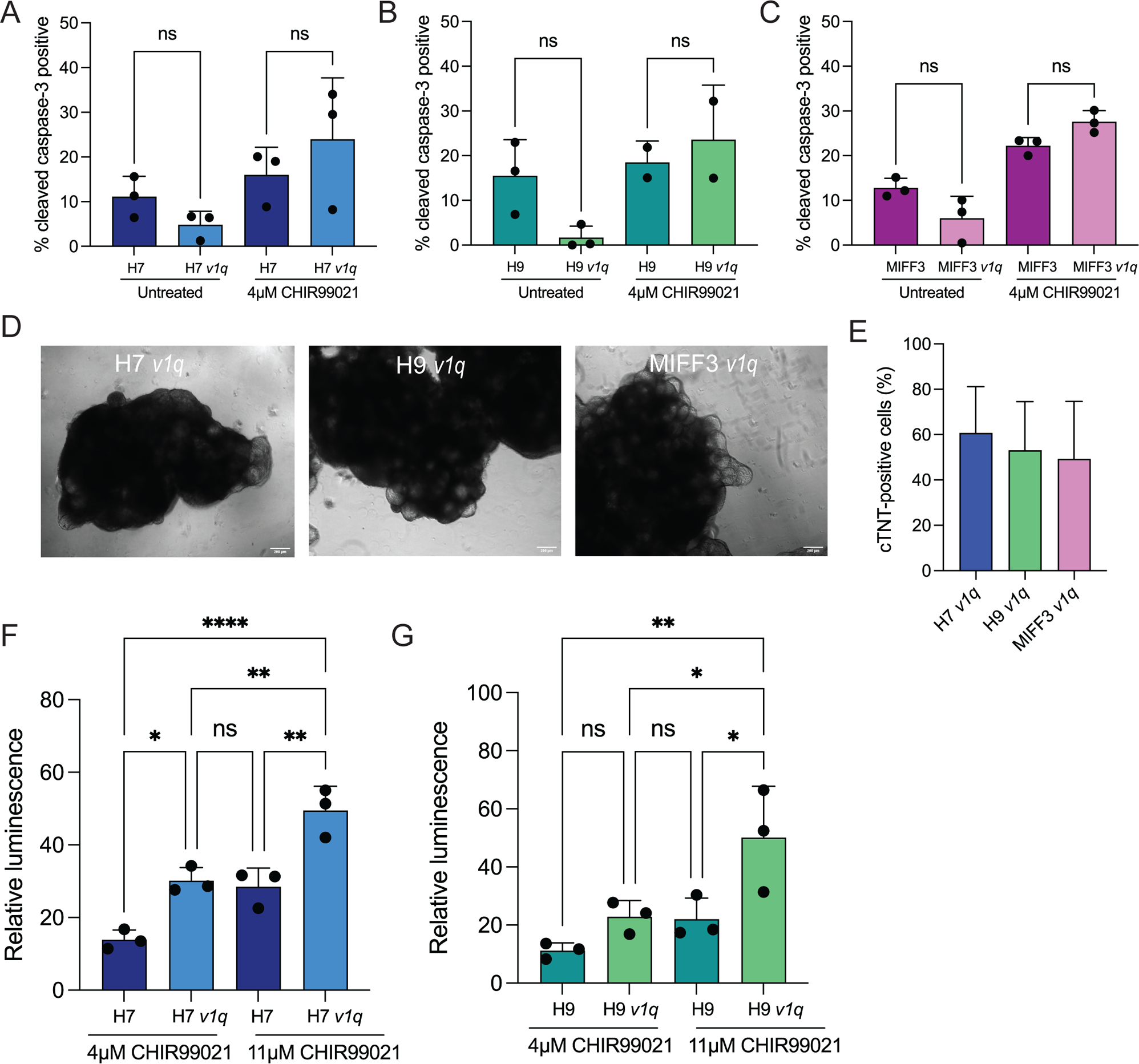
Hyperresponsiveness to Wnt activation impairs differentiation of variant 1q hPSCs to cardiomyocytes. (A) Reducing the CHIR99021 concentration lowers the amount of apoptosis in variant 1q lines. Percentage of caspase-3 positive H7 and H7 *v1q* cells grown with or without 4µM CHIR99021 for 24h. (B) Percentage of caspase-3 positive H9 and H9 *v1q* cells grown with or without 4µM CHIR99021 for 24h. (C) Percentage of caspase-3 positive MIFF3 and MIFF3 *v1q* cells grown with or without 4µM CHIR99021 for 24h. (D) Representative images of *v1q* cells across three different genetic backgrounds (H7, H9 and MIFF3) at day 8 of differentiation on microcarriers using a reduced concentration of CHIR99021. Scale bar: 200 µm. (E) Reducing the CHIR99021 concentration increases the differentiation efficiency of variant 1q lines. Percentage of cardiac troponin (cTNT) -positive cells for the three WT lines treated with either 1µM CHIR99021 (H7 *v1q*) or 4µM CHIR99021 (H9 *v1q* and MIFF3 *v1q*) in the microcarrier cardiomyocyte differentiation protocol (day 8). H7*v1q* n=3, H9 *v1q* and MIFF3 *v1q* n=2. (F) Wnt activation measured as relative luminescence of TOPflash versus FOPflash luminescent signal in cells treated with 4µM and 11µM CHIR99021 (H7 and H7 *v1q*) (G) Wnt activation measured as relative luminescence of TOPflash versus FOPflash luminescent signal in cells treated with 4µM and 11µM CHIR99021 (H9 and H9 *v1q*) Results are the mean of three independent experiments ± SD, unless stated otherwise. ns=non-significant; *p<0.05; **p<0.01; ****p<0.0001, One-way ANOVA with Tukey’s multiple comparison test.

To test our assumption that differentiation in *v1q* lines upon treatment with lower doses of CHIR99021 was due to achieving just right levels of Wnt activation, we employed the TOP-Flash Wnt reporter assay^22^ on variants treated with optimised levels of CHIR99021 (4µM). This analysis confirmed that the levels of Wnt activation with lower doses of 4µM CHIR99021 in *v1q* sublines were equivalent to the Wnt activation in WT cells with 11µM CHIR99021 (**Figure 3F,G**). Together, our data shows that the excessive cell death of variant cells is caused by hyperactivation of Wnt and reveals that the levels of Wnt must be tightly regulated to achieve efficient cardiomyocyte differentiation.

### Transcriptomic and phenotypic analysis of cardiomyocytes derived from WT and *v1q* hPSCs

We next asked whether the cardiomyocytes derived from *v1q* cells are phenotypically equivalent to those produced from WT cells. To address this, we first performed genome-wide transcriptome profiling by RNA-seq in WT and *v1q* H7 cells at the undifferentiated (d0) and cardiomyocyte-differentiated state (d8). To achieve efficient differentiation, here we used differentiation protocols specifically optimised for each of these sublines, i.e. using 11µM CHIR99021 for WT and 1µM for *v1q* H7 cells. Principal component analysis showed that while *v1q* and WT cells are transcriptionally relatively similar to each other at d0, they diverge significantly at the cardiomyocyte state (d8). Specifically, *v1q* cells are deficient in genes associated with normal cardiomyocyte differentiation and remain clustered with hPSC-state (d0) based on PC1 component (**Figure S2A**).

To better understand the transcriptional changes, we computed differentially expressed genes at each stage of differentiation and detected n=591 and n=8,876 high confidence differentially expressed genes between *v1q* and WT cells at d0 and d8, respectively (log2FC > 1.5, and FDR <0.05, **Figure S2B** and **Table S1**). The majority of differentially expressed genes at d0 were upregulated in *v1q* cells (n=495) whereas a more comparable number of up-(n=3,352) and down-regulated (n=5,524) genes were detected in *v1q* cells at d8 of differentiation. Furthermore, the number of differentially expressed genes were distributed across multiple chromosomes at d8 of differentiation ruling out a major bias in transcriptional profiles towards chr1 loci (**Figure S2C**). GO-term enrichment analysis revealed that genes downregulated in *v1q* cells at cardiomyocyte stage (d8) are mainly associated with cardiomyocyte differentiation including genes involved in extracellular matrix interaction, collagen and fibre formation, and muscle contraction. In contrast, genes up-regulated in *v1q* cells at d8 were predominantly involved in cell cycle pathways (**Figure S2D**). Wnt signalling was also one of the pathways identified as altered in *v1q* cells at d8 of differentiation (**Table S1**). Altogether, these data indicated that multiple genes are dysregulated in variant hPSCs in undifferentiated state but the transcriptomes of WT and *v1q* lines become profoundly different upon differentiation.

In line with marked transcriptional changes, the phenotypic assessment of α-actinin-stained cells after eight days of differentiation also showed overt differences between WT- and *v1q*-derived differentiated cells (**Figure 4A,B**). Notably, some of the differences, such as the cell size, appeared cell line-specific, with MIFF3 *v1q* α-actinin positive cells appearing significantly smaller and H9 *v1q* cells significantly larger than WT counterparts (**Figure 4C**). Conversely, a common feature across cardiomyocytes derived from all three *v1q* lines was a significantly shortened average branch length of α-actinin positive fibres (**Figure 4D**). The total length of α-actinin fibers (**Figure 4E**) and the number of cardiac fibre junctions per cell (**Figure 4F**) were overall similar between WT- and *v1q*-derived cardiomyocytes. Collectively, this data revealed that although variant cells are capable of differentiation when Wnt activation is finely tuned, the resulting cardiomyocytes deviate significantly from their wild-type counterparts.

**Figure 4.**
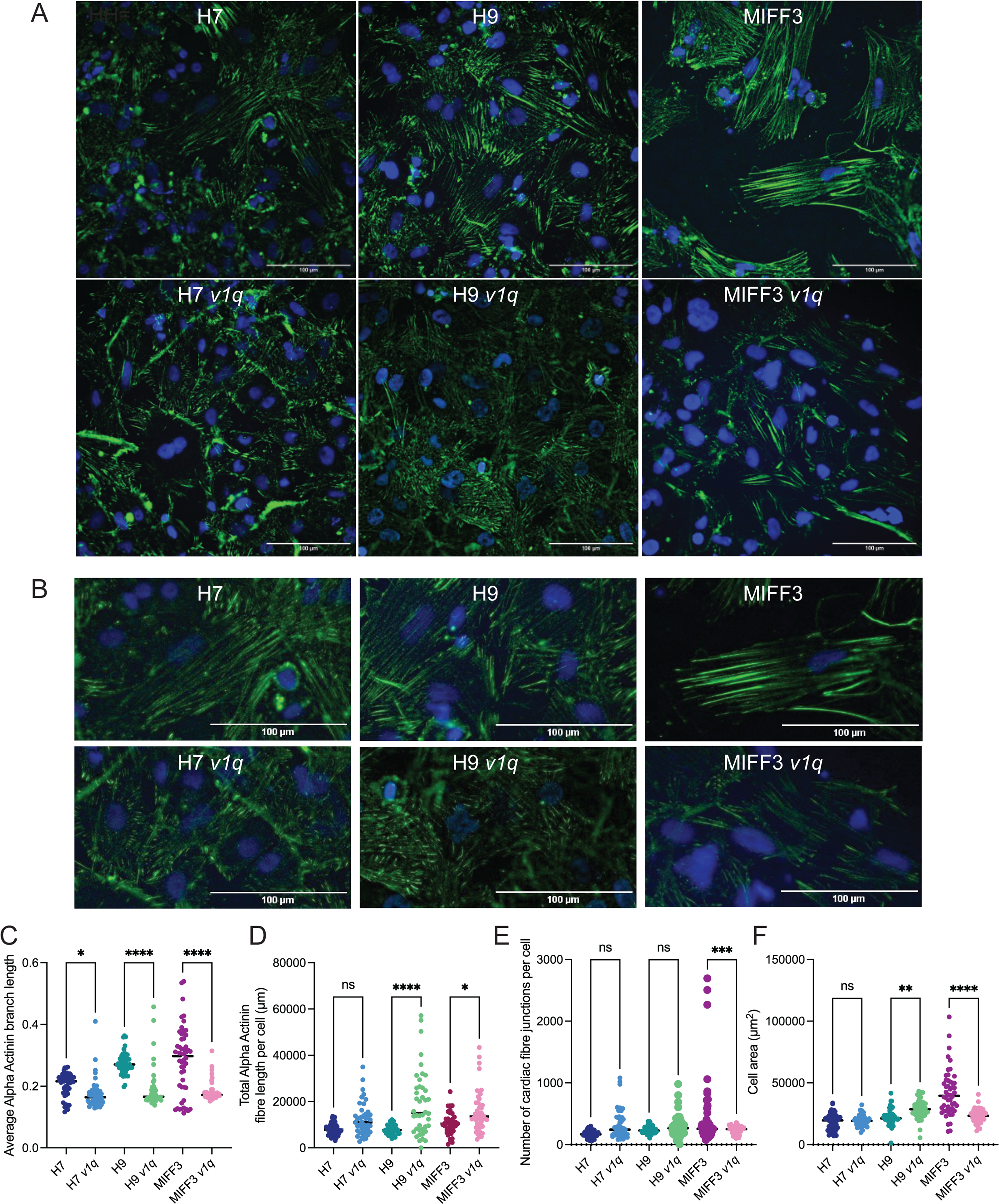
Cardiomyocytes generated by differentiating variant 1q hPSCs are phenotypically different from their WT counterparts. (A) Representative images of WT and v1q cardiomyocytes across three different genetic backgrounds (H7, H9 and MIFF3) generated by differentiating cells on microcarriers for 8 days using an optimised CHIR99021 concentration for v1q lines. After 8 days of differentiation on microcarriers, cells were replated into dishes, and after 2 days fixed and stained with α-actinin antibody (green). Nuclei were counterstained with Hoechst 33342 (blue). Scale bar: 100µm. (B) The zoomed-in images of α-actinin staining from Figure 4A. Scale bar: 100µm. (C) Cell size of α-actinin positive cells. 50 fields of α-actinin positive cells per cell line were imaged and quantified. n.s. non-significant; **p<0.01; ****p<0.0001; One-way Anova for individual pairs of WT and *v1q* pairs of sublines. (D) Average α-actinin positive branch length quantified from images of α-actinin positive WT and *v1q* cells. 50 fields of α-actinin positive cells per cell line were imaged and quantified. *p<0.05; ****p<0.0001; One-way Anova for individual pairs of WT and *v1q* pairs of sublines. (E) Total length of α-actinin positive fibres per cell quantified from images of α-actinin positive WT and *v1q* cells. 50 fields of α-actinin positive cells per cell line were imaged and quantified. n.s. non-significant; *p<0.05; One-way Anova for individual pairs of WT and *v1q* pairs of sublines. (F) The number of junctions between α-actinin positive fibres per cell quantified from images of α-actinin positive WT and *v1q* cells. 50 fields of α-actinin positive cells per cell line were imaged and quantified. n.s. non-significant; ***p<0.001; One-way Anova for individual pairs of WT and *v1q* pairs of sublines.

## Discussion

The inefficiency of hPSC differentiation and/or altered functional characteristics of differentiated cells due to the presence of genetic changes in the starting hPSC population stand to seriously jeopardise the use of hPSC-derived cells in research or therapy. In this study, by using three independent pairs of WT and variant lines with a gain of chromosome 1q in two different differentiation protocols, we established that the presence of the chromosome 1q gain affects the differentiation of hPSC to cardiomyocytes. We further demonstrated that this defect is due to an altered sensitivity of variant cells to Wnt activation. First, we showed that the same dose of CHIR99021 that triggers differentiation in WT cells causes apoptosis in *v1q* hPSCs. We confirmed that this effect was due to Wnt activation as we could phenocopy it by using the natural Wnt ligand, Wnt3a. Second, we demonstrated that this differential response to Wnt agonists is likely because variants are hyperresponsive to Wnt activation, as the same dose of CHIR99021 caused a much quicker and greater upregulation of Wnt signalling, as measured by TOPFlash assay. Finally, by decreasing the levels of CHIR99021, we were able to overcome the apparent block in differentiation and achieve cardiomyocyte differentiation of *v1q* hPSCs. This latter result demonstrated that *v1q* hPSCs are not inherently recalcitrant to differentiation but rather, genetic changes that variant hPSCs harbour affect their response to differentiation cues.

While we were able to alleviate differences in the propensity for differentiation of variant cells by fine-tuning their signalling environment, the resulting differentiated cells significantly differed from their WT counterparts. For example, apart from an increase in the expression of genes harboured within the amplified chromosomal region, transcriptome analysis highlighted global transcriptional changes. Many pathways were significantly dysregulated as a result, including the Wnt pathway, although we do not yet know the precise mechanism by which the gain of chromosome 1q influences the capacity of Wnt activation. Further phenotypic differences were evident from the immunocytochemistry analysis of alpha actinin-positive cardiac fibre length and connectivity, which represent a hallmark of cardiomyocyte maturity and functionality^23^. Although we noted some variability in the cardiomyocyte phenotype of WT lines, *v1q* cells consistently differed from their paired WT counterparts. All *v1q* lines examined displayed shorter fibre lengths, suggesting a more immature phenotype. Future work should address the extent of functional consequences of recurrent chromosome 1q gain on hPSC-derived cardiomyocytes. Of note, a parallel study by Brandao et al. (2023) showed enhanced contractility of a *v1q* line in a 3D cardiac model, thus demonstrating that recurrent genetic changes in hPSC can have profound consequences for functional behaviour of their differentiated derivatives.

There are at least a couple of ways in which *v1q* hPSCs can undermine the effectiveness of differentiation protocols for research or cell therapy. First, some of the variants investigated in our study, such as MIFF3 *v1q*^15^, had a relatively small amplification of chromosome 1q, which appeared below the resolution of G-banding^15^, but nonetheless affected the differentiation efficiency of hPSCs. The relatively small size of this CNV limits the detection of variant cells^8^, making them more likely to go unnoticed in cultures. Second, even if harbouring larger karyotypic abnormalities, variant cells may be present at a low proportion of cells in culture. Low level of mosaicism also presents detection challenges of variants^8^. As a corollary, the contamination of cultures with variant cells may impact the differentiation efficiency of protocols used. Indeed, given the striking effects of variants on the differentiation efficiency of hPSCs^24^, it seems plausible that at least some of the widely reported batch-to-batch variations in differentiation performance could be due to the presence of culture-acquired genetic changes arising in hPSCs and consequently taking over the cultures. On the other hand, our observation that *v1q* hPSCs are more prone to apoptosis upon Wnt activation may offer a practical solution for the elimination of variant cells during expansion. Specifically, titrating the levels of Wnt may allow concomitant differentiation of WT cells and elimination of *v1q* from mosaic cultures.

In summary, our work demonstrates that recurrently gained genetic variants in hPSCs alter the differentiation capacity of hPSCs to cardiomyocytes. We identified enhanced susceptibility to Wnt activation as the key reason for this difference. Given the commonality of recurrent genetic changes in hPSC cultures, the presence of variants may be the prime reason for issues with reproducibility and robustness of the hPSC differentiation protocols. Our data suggests that copy number changes affecting the Wnt pathway activation may be implicated in aberrations associated with cardiomyocyte differentiation during development. Finally, our study demonstrates a paradigm approach for investigating the impact of genetic variants on hPSC differentiation and behaviour of other specialized cell types, relevant for disease modelling and cell therapy.

## EXPERIMENTAL PROCEDURES

### Resource availability

#### Corresponding authors

Further information and requests for resources and reagents should be directed to and will be fulfilled by the corresponding author, Ivana Barbaric.

#### Materials availability

This study did not generate new unique reagents.

#### Human pluripotent stem cell (hPSC) lines

Wild-type hPSCs used in this study were H7 (WA07)^1^, H9 (WA09)^1^ and MIFF-3^17^. Wild-type sublines were karyotypically normal (based on at least 20 metaphases analysed by G-banding of cell banks prior to experiments). Cells with a gain of chromosome 1q used in this study were: H7 *v1q*^12^ [46,XX,dup(1)(q21q42)] (20 metaphases analysed), H9 *v1q*^15^ [46,XX,der(21)t(1;21)(q21;p11)] (20 metaphases analysed), MIFF-3 *v1q*^15^ [46,XY] (20 metaphases analysed). While MIFF-3 *v1q* seemed karyotypically normal by G-banding, a duplication of q32.1 on chromosome 1 was detected by qPCR and confirmed by SNParray analysis^15^. All lines were characterised at the time of banking and cells from the bank were used no more than 10 passages from defrosting.

#### Cell culture

For routine culture, hPSCs were grown in flasks coated with Geltrex LDEV-Free Reduced Growth Factor Basement Membrane Matrix (Gibco, A1413202) diluted at 1:100 in DMEM/F12 (Sigma-Aldrich, D6421). The medium used for hPSC maintenance was a modified E8 medium^25^, prepared in house. Cells were routinely passaged utilising ReLeSR (STEMCELL Technologies, 100-0484) according to manufacturer’s instructions.

#### Cardiomyocyte differentiation

2D cardiomyocyte differentiations were performed utilising either the STEMdiff ventricular cardiomyocyte differentiation kit (STEMCELL Technologies, 05010) or using a microcarrier differentiation protocol adapted from previous studies^18,19^.

#### Luciferase reporter assay

For the Wnt reporter assay, 2.5 million hPSCs were transfected with either 400ng WRE plasmid with active TCF binding sites (Addgene, 12456) or MRE plasmid with mutated TCF binding sites (Addgene, 12457), together with 100 ng of the Renilla luciferase control plasmid (Addgene, 118016). Transfections were performed using a Neon transfection kit (Thermo-Fisher, MPK10025). Luciferase activity was measured using a Dual–Luciferase Reporter Assay System (Promega, E1500).

#### Immunocytochemistry

Immunocytochemistry was performed as previously described^12^. Briefly, cells were fixed with 4% paraformaldehyde for 10 minutes at room temperature and then blocked and permeabilised with 10% foetal calf serum (FCS) supplemented with 0.2% Triton X-100 in PBS for 10 minutes RT or 1 hour at 4°C. Cells were incubated with a primary antibody in 10% FCS at 4°C overnight. After three washes with PBS, cells were incubated with a secondary antibody in 10% FCS for 1 hour. The list of antibodies is provided in Supplemental Information.

#### Cleaved caspase-3 assay

Cells were harvested from a 12 well plate, with cells in medium first pelleted with adherent cells harvested with TrypLE express (Gibco, 12604021). Cell pellets were then stained utilising protocol seen above under Immunocytochemistry with cells stained with cleaved caspase-3 rabbit polyclonal primary antibody for 1 hour (Cell Signalling Technology, 9661). Percentages of cleaved caspase-3 positive cells were determined utilising Flow cytometry, BD FACSJazz, against an unstained control.

#### RNAseq

RNAseq was performed using Genewiz RNAseq service, with triplicate samples of day 0 hPSCs and day 8 3D differentiated cardiomyocytes undergoing RNAseq. FASTQ reads were mapped against hg19 reference genome using *STAR*^26^ and default setting. Mapped reads were assigned to RNAs using HTseq-count^27^ with the following settings: -m union -s reverse -t exon. Samples were further analyzed by DESeq2^28^ (bioconductor.org) for differential gene expression analysis and with the following cut-off: log2 reads > 5 in at least one sample, FDR<0.05, log2 FC > 1.5). Graphs were generated in R version 4.3.1 on Ubuntu 16.04.5 LTS. GO-term enrichment analysis was performed using *gprofiler* (https://biit.cs.ut.ee/gprofiler/gost).

## Author Contributions

TW, SO, YA and IB conceived and designed experiments. TW, CP, DS, OL, JR, AL, YA and IB performed the experiments and/or analysed the data. IB wrote the manuscript with input from all authors.

## Acknowledgements

This work was supported by the UK Regenerative Medicine Platform (MR/R015724/1), the Medical Research Council (MR/X000028/1, MR/X007979/1, MR/N013840/1) and BBSRC DTP (BB/M011151/1, BB/T007222/1) studentships. We thank Sheffield Genetics Diagnostics Service for cytogenetic analyses. Imaging work was performed at the Wolfson Light Microscopy Facility at the University of Sheffield, using the ZEISS LSM880 AiryScan microscope and Nikon W1 spinning disk confocal (BB/V019368/1).

## Supplemental Information

### Supplemental Figures

**Figure S1, related to Figure 1 and Figure 2.**
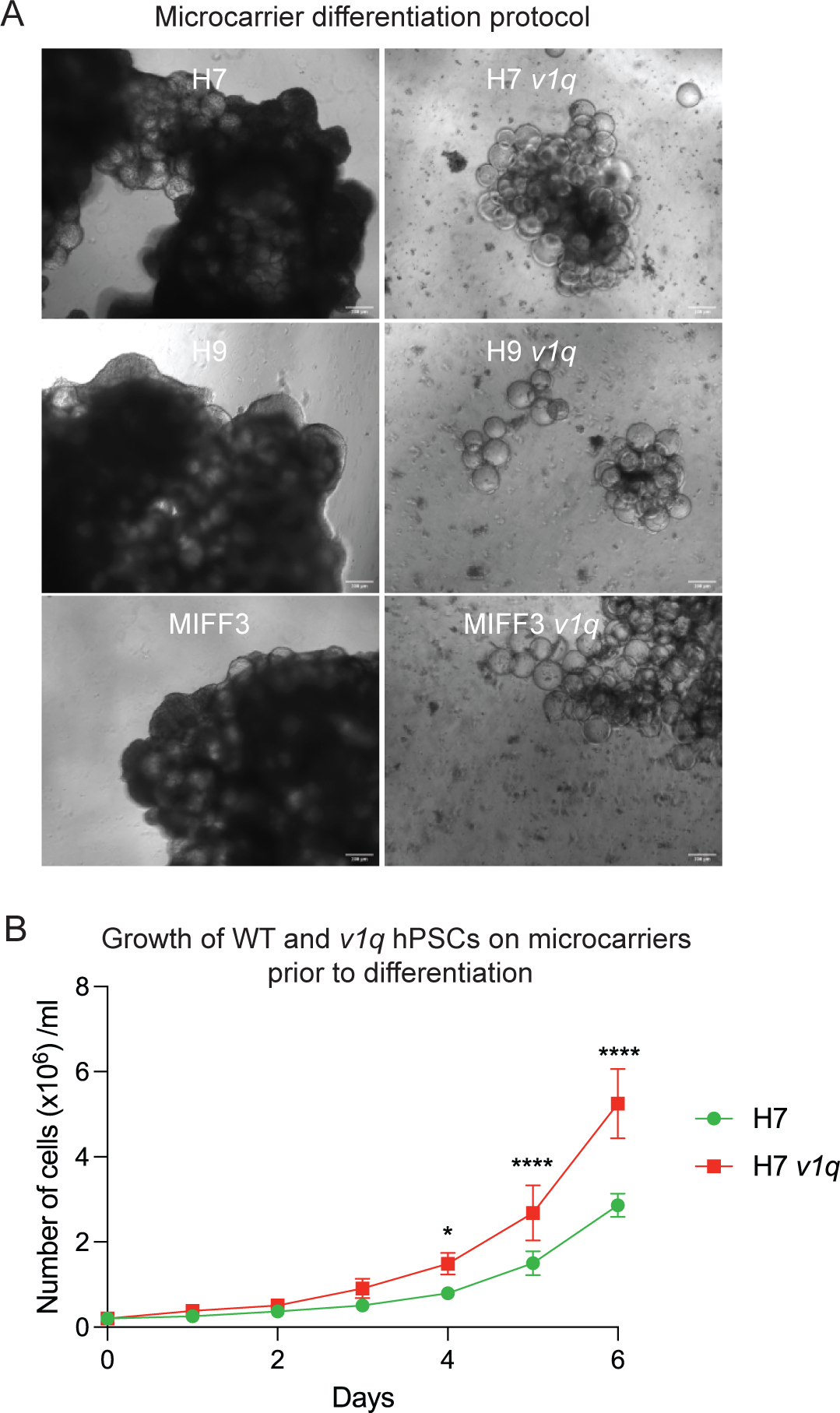
Variant hPSCs with a gain of chromosome 1q die excessively upon differentiation on microcarriers, although initial growth of variant cells on microcarriers is unaffected. (A) Representative images of WT and *v1q* cells across three different genetic backgrounds (H7, H9 and MIFF3) at day 8 of differentiation on microcarriers using the 3D differentiation kit. Scale bar: 200µm. (B) Growth curves of H7 wild-type and H7 *v1q* hPSCs grown on microcarriers in hPSC medium. Results are the mean of three independent experiments ± SD. *p<0.05; ****p<0.0001, Two-way ANOVA with Tukey’s multiple comparison test.

**Figure S2, related to Figure 4.**
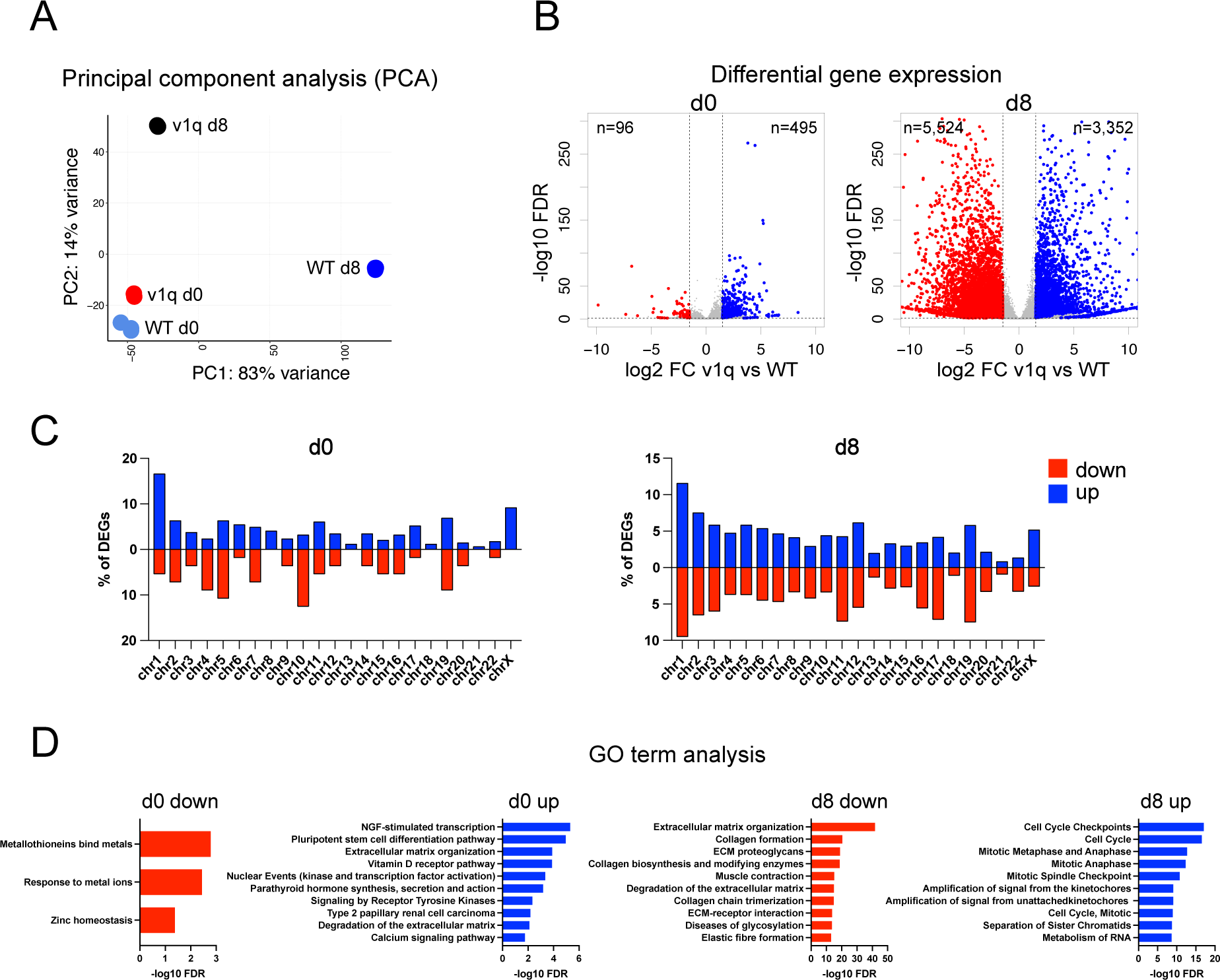
Cardiomyocytes generated by differentiating *v1q* hPSCs are transcriptionally different from their WT counterparts. (A) Transcriptional profiling of WT and *v1q* cells in undifferentiated state (d0) and upon differentiation to cardiomyocytes (d8). Principal component analysis of WT and *v1q* H7 cells in undifferentiated state (d0) and at day 8 of the microcarrier cardiomyocyte differentiation protocol (d8). Three biological replicates were used per experimental group. (B) Differentially expressed genes corresponding to ***panel-A***. Blues and red dots represent up- and down-regulated genes respectively, with a log2 fold change >1.5, and FDR < 0.05. n represents the number of differentially expressed genes. (C) Barplot represents the distribution of differentially expressed genes (DEGs) across different chromosomes. Blue and red bars represent up- and down-regulated genes in *v1q* vs WT cells respectively (corresponding to ***panel-B***). (D) GO term enrichment analysis for KEG/REACT/WP pathways for differentially expressed genes at d0 and d8 of differentiation. Blue and red bars represents gene up- and down-regulated in *v1q* vs WT cells respectively (corresponding to ***panel-B***). Only the top 10 enriched GO-terms are shown for each gene list.

### Supplemental Videos

**Video S1.**

Panel of H7, H9 and MIFF3 functional beating cardiomyocytes at day 10 of differentiation using the STEMdiff ventricular cardiomyocyte differentiation kit (STEMCELL Technologies). Videos were taken at 24 frames per second.

**Video S2.**

Panel of H7 *v1q*, H9 *v1q* and MIFF3 *v1q* at day 10 of differentiation using the STEMdiff ventricular cardiomyocyte differentiation kit (STEMCELL Technologies). Videos were taken at 24 frames per second.

**Video S3.**

Panel of H7, H9 and MIFF3 functional beating cardiomyocytes aggregated on Cytodex-1 microcarriers at day 8 of 3D cardiomyocyte differentiation. Videos were taken at 10 frames per second.

**Video S4.**

Panel of H7 *v1q*, H9 *v1q* and MIFF3 *v1q* functional beating cardiomyocytes aggregated on Cytodex-1 microcarriers at day 8 of 3D cardiomyocyte differentiation protocol utilising an optimised CHIR99021 concentration for each variant line. Videos were taken at 10 frames per second.

### Supplemental Experimental Procedures

#### Karyotyping

Karyotyping by G-banding was performed by the Sheffield Diagnostic Genetics Service (https://www.sheffieldchildrens.nhs.uk/sdgs/), as previously described^1^. Karyographs were analysed by a Clinical Scientist at the Sheffield Diagnostic Genetics Service. An average of 20 metaphases were analysed by G-banding per sample.

#### Modified E8 medium for hPSC maintenance

Modified E8 medium^2^ was prepared by supplementing DMEM/F12 (Sigma-Aldrich, D6421) with 14 µg/l sodium selenium (Sigma-Aldrich, S5261), 19.4 mg/l insulin (Thermo Fisher Scientific, A11382IJ), 1383 mg/l NaHCO_3_ (Sigma-Aldrich, S5761), 10.7 mg/l transferrin (Sigma-Aldrich, T0665), 10 ml/l Glutamax (Thermo Fisher Scientific, 35050038), 40µg/l FGF2-3 (Qkine, Qk053) and 2 µg/l TGFβ1 (Peprotech, 100-21).

#### RPMI 1640 medium for hPSC differentiation

RPMI 1640 basal differentiation media was prepared in house as previously described^3^. Briefly, RPMI 1640 basal media powder (Thermo-Fisher, 31800105) was dissolved in culture grade distilled water. This basal media was supplemented with sodium bicarbonate, L-Ascorbic acid (Thermo-Fisher, A15613.36) and B27 minus insulin (Gibco, 15285074).

#### 2D cardiomyocyte differentiation

2D cardiomyocyte differentiations were performed utilising the STEMdiff ventricular cardiomyocyte differentiation kit (STEMCELL Technologies, 05010), according to the manufacturer’s instructions. Briefly, hPSCs were seeded at a density of 150,000 cells/cm^2^ and cultured in mTeSR Plus (STEMCELL Technologies) for 48 hours, with the media in the first 24 hours containing 10µM Y-27632. Medium A was then used for the next 48 hours, followed by medium B for another 48 hours and subsequently medium C for 96 hours. After this period, cardiomyocyte maintenance medium was used.

#### Microcarrier cell culture and 3D differentiation

hPSCs were dissociated from monolayer flasks and resuspended at a 1 million cells/ml in modified E8 media supplemented with 10µM Y-27632. Geltrex-coated Cytodex-1 (Cytiva, 17044801) microcarriers were centrifuged at 2500 rpm for 5 minutes to form pellet. Supernatant was aspirated, with close care taken to avoid aspiration of microcarrier pellet. The microcarrier pellet was resuspended in modified E8 media supplemented with 10µM Y-27632 and 4 ml of suspended microcarriers were added to each well of 6 well ultra-low attachment plate (Corning, CLS3471-24 EA). To each well, 1 million hPSCs were added. Plates were then placed at 37°C on orbital shaker for 2 hours to allow aggregation. After this two-hour period, visible aggregates were typically seen, and plates were moved to static 37°C incubator overnight. Medium was refreshed every 24 hours.

The cardiomyocyte differentiation protocol was typically initiated 5-7 days after plating the cells on microcarriers. Aggregated cells were counted in order to determine the number of aggregates required per well. Microcarrier aggregates were taken from individual wells and collated in 50 ml falcon tube and allowed to settle in the base. From the 50 ml falcon tube, all media was aspirated, and cells were washed with 20 ml of PBS. Once more cells were allowed to settle, and the PBS was aspirated. A secondary wash was performed utilizing 20 ml of RPMI 1640 basal differentiation media. This also was subsequently aspirated, and the wash was repeated. Lastly, according to cell numbers, RPMI with the addition of CHIR-99021 (Tocris, 4423) was added to the tube. The cells were then ready to be added to a 6-well ultra-low attachment plate, with 5 ml of suspended cells at a cellular density of 1 million cells per well. This was performed using a 25ml serological pipette, with gentle resuspension occurring after every well to ensure the microcarrier aggregates remained in suspension. Once the cells were replated, the plates were transferred back to 37°C for 18 hours.

After 18 hours, the RPMI 1640 supplemented with CHIR-99021 was aspirated and cells were washed 3 times with 3 ml of basal differentiation media. The final wash was replaced with 5 ml of RPMI 1640, and the cells were placed on an orbital shaker at 37°C for 48 hours. At day 3 of the differentiation protocol, cells were treated with 5µM IWR-1 (Merck 10 161-25mg). Once more, media was aspirated, and the microcarrier aggregates underwent three washes with RPMI 1640. The final wash replaced the media with 5 ml of RPMI 1640 supplemented with 5µM IWR-1. The plates were replaced at 37°C for a further 48 hours.

This supplementation of the differentiation media remained for 48 hours, at which point the cellular treatment with small molecules was complete. The IWR-1 containing media was aspirated and three 3ml RPMI washes occurred. This media replacement with basal RPMI 1640 differentiation media was repeated every 48 hours until the cardiomyocyte phenotype was seen, with the cells remaining on an orbital shaker from day 5 onwards.

#### Immunocytochemistry

Cells were fixed with 4% PFA for 10-15 minutes at room temperature and then washed with PBS three times. Cells were subsequently blocked and permeabilized with PBS supplemented with 1% BSA, 0.3% Triton X-100 and 0.2% Tween-20 for 10 minutes at room temperature or 1 hour at 4°C. Cells were washed three times with PBS and then incubated with primary antibody in PBS supplemented with 10% foetal calf serum for 1 hour at room temperature or overnight at 4°C. Cells were subsequently washed twice with PBS before incubating with secondary antibody. Secondary antibodies were also resuspended in blocking buffer and Hoechst 33342 (ThermoFisher, H3570) was added at 1:1000 dilution. Cells were incubated with secondary antibody/ Hoechst 33342 for one hour at 4°C. Cells were again washed three times with PBS before imaging with an InCell Analyzer 2200 (GE Healthcare). Control samples-only stained with secondary antibody were utilized as a baseline control to measure fluorescence intensity.

Microcarrier aggregates did not undergo permeabilization, before staining, with 0.3% Triton X-100 not added to blocking buffer. Post blocking aggregates were treated with Hoechst 33342 (ThermoFisher, H3570) at 1:1000 dilution alongside phalloidin, as stated below.

Images were obtained utilising GE healthcare IN Cell 2200 analyser, Zeiss LSM880 Airyscan confocal or W1 Spinning Disk confocal microscopes.

#### Flow cytometry

Cells were dissociated from both monolayer and microcarrier cultures utilizing TrypLE express (Gibco, 12604021), with a 5-minute treatment at RT for hPSC cultures and 1 hour at RT for cardiomyocyte cultures. Cells subsequently underwent staining protocol as seen above with Immunocytochemistry section, with cells prepared for flow cytometry in 15 ml falcons. List of antibodies utilised in this study can be seen below.

Flow cytometry was performed utilising BD FACSJazz flow cytometer. Flow cytometer was calibrated before every use using BD Sphero Rainbow calibration particles, 8 peaks (Fisher scientific, 15859678). Unstained control samples were performed to set baseline of cellular autofluorescence, with gate set at highest 5% of intensity as a positive sample.

#### Antibodies for immunofluorescence and flow cytometry

| **Antibody** | **Type** | **Dilution used** | **Supplier** |
| --- | --- | --- | --- |
| Myosin 4 | Mouse Monoclonal | 1:100 | Abcam, ab37484 |
| Cardiac Troponin | Rabbit Monoclonal | 1:200 | Thermo-Fisher,<br>MA5-12960 |
| Cleaved caspase-3 | Rabbit Polyclonal | 1:400 | Cell Signalling<br>Technology, 9661 |
| Alpha actinin | Mouse Monoclonal | 1:200 | Abcam, ab18061 |
| Myosin light chain<br>2V | Rabbit Monoclonal | 1:200 | Proteintech,<br>10906-1-AP |
| Phalloidin | Conjugated 647<br>polyclonal | 1:50 | Thermo-Fisher<br>A30107 |
| Mouse primary<br>antibodies | Goat anti-mouse IgG<br>Alexa 488 | 1:1000 | Invitrogen,<br>A11001 |
| Mouse primary<br>antibodies | Goat anti-mouse IgG<br>Alexa 594 | 1:1000 | Invitrogen,<br>A11005 |
| Rabbit primary<br>antibodies | Goat anti-rabbit IgG<br>Alexa 488 | 1:1000 | Invitrogen,<br>A11008 |
| Rabbit primary<br>antibodies | Goat anti-rabbit IgG<br>Alexa 594 | 1:1000 | Invitrogen,<br>A11012 |

#### Image quantification

Image quantification was performed using Fiji^4^ pipelines, with Tiff image files converted to a binary mask. Binary images skeletons were then summarised giving readouts of average branch length, total branch length and number of branch junctions. Cell numbers and size were obtained using Cell profiler^5^, with Hoechst 33342 and α-actinin staining utilised as primary and secondary objects. For number of cell junctions per cell readouts the total number of branch junctions were normalised using Cell profiler cell counts.

#### RNA-seq and data analysis

RNA-seq in this study was performed using Genewiz RNA-seq service. RNA-seq libraries were sequenced on a HiSeq-2000 Illumina sequencer. FASTQ reads were mapped against hg19 reference genome using *STAR*^6^ and default setting. Mapped reads were assigned to RNAs using HTseq-count^7^ with the following settings: -m union -s reverse -t exon. Samples were further analyzed by DESeq2^8^ (bioconductor.org) for differential gene expression analysis and with the following cut-off: log2 reads > 5 in at least one sample, FDR<0.05, log2 FC > 1.5). Graphs were generated in R version 4.3.1 on Ubuntu 16.04.5 LTS. GO-term enrichment analysis was performed using *gprofiler* (https://biit.cs.ut.ee/gprofiler/gost)

